# PLK-1 tethered on BUB-1 directs CDC-20 kinetochore recruitment to ensure timely embryonic mitoses

**DOI:** 10.1101/2022.10.07.511323

**Authors:** Jack Houston, Midori Ohta, J. Sebastián Gómez-Cavazos, Amar Deep, Kevin D. Corbett, Karen Oegema, Pablo Lara-Gonzalez, Taekyung Kim, Arshad Desai

## Abstract

During mitosis chromosomes assemble kinetochores in order to dynamically couple with spindle microtubules (Cheeseman, 2014; Musacchio & Desai, 2017). Kinetochores also function as signaling hubs directing mitotic progression by recruiting and controlling the fate of the Anaphase Promoting Complex/Cyclosome (APC/C) activator CDC-20 (Lara-Gonzalez et al., 2017; Lara-Gonzalez, Pines, et al., 2021; Musacchio, 2015). Kinetochores either incorporate CDC-20 into checkpoint complexes that inhibit the APC/C or dephosphorylate CDC-20, which allows it to interact with and activate the APC/C (Kim et al., 2017; Lara-Gonzalez et al., 2017). The importance of these two CDC-20 fates likely depends on biological context. In somatic cells the major mechanism controlling mitotic progression is the spindle checkpoint. By contrast, progression through mitosis during the cell cycles of early embryos is largely checkpoint-independent (Clute & Masui, 1995; Duro & Nilsson, 2021; Gerhart et al., 1984; Zhang et al., 2015). Here, by manipulating CDC-20 phosphorylation status, we show that CDC-20 phosphoregulation controls mitotic duration in the *C. elegans* embryo and defines a checkpoint-independent temporal mitotic optimum for robust embryogenesis. Flux of CDC-20 through kinetochores for local dephosphorylation requires an ABBA motif on BUB-1 that directly interfaces with the structured WD40 domain of CDC-20 (Di Fiore et al., 2015; Diaz-Martinez et al., 2015; He et al., 2013; Kim et al., 2017). We show that a conserved “STP” motif in BUB-1 that docks the mitotic kinase PLK-1 (Qi et al., 2006) is also necessary to recruit CDC-20 to kinetochores and for timely mitotic progression. The kinase activity of PLK-1 is required for CDC-20 to localize to kinetochores and targets a site within the CDC-20-binding ABBA motif of BUB-1; phosphorylation of this site promotes BUB-1–CDC-20 interaction and mitotic progression. Thus, the BUB-1-bound pool of PLK-1 ensures timely mitosis during embryonic cell cycles by promoting CDC-20 recruitment to the vicinity of kinetochore-localized phosphatase activity.

## RESULTS

### CDC-20 phosphoregulation optimizes mitotic duration to ensure robust embryogenesis

In the early *C. elegans* embryo at 20°C the alignment and onset of separation of all 12 chromosomes is achieved during the ~180-200 second interval that follows nuclear envelope breakdown (NEBD). Eliminating spindle checkpoint signaling does not alter this timing (Essex et al., 2009; Kim et al., 2017), indicating that tight control of mitotic duration during these rapid embryonic divisions is achieved through a checkpoint-independent mechanism.

In prior work, we found that phosphoregulation of the APC/C activator CDC-20 is an important regulatory mechanism that controls mitotic duration in the early *C. elegans* embryo (Kim et al., 2017). Phosphorylation of Cdc20 by Cdk1/2 on sites adjacent to the C-box, a critical APC/C binding motif in the N-terminus, prevents Cdc20 from binding and activating the APC/C *in vitro* and in cells (Alfieri et al., 2017; Kramer et al., 2000; Labit et al., 2012; Yudkovsky et al., 2000) (**Fig. 1A)**. Dephosphorylation of Cdc20 by protein phosphatases is critical to promote its binding to the APC/C and thereby anaphase onset (Bancroft et al., 2020; Hein et al., 2021; Hein & Nilsson, 2016; Shevah-Sitry et al., 2022). In *C. elegans* preventing the inhibitory phosphorylation of CDC-20 by mutating 3 potential Cdk sites in its N-terminus to alanine significantly accelerated mitotic progression, shortening the NEBD-to-anaphase interval to ~120 seconds, indicating premature APC/C activation in the absence of CDC-20 phosphorylation (Kim et al., 2017). The most functionally significant of these 3 sites is Thr32, which is positioned adjacent to the critical APC/C binding C-box. A single Thr32Ala mutation in CDC-20 accelerates mitosis to a similar extent as the triple alanine mutant (**Fig. 1B-D**; (Kim et al., 2017)). To determine whether persistent CDC-20 phosphorylation has the opposite effect, we had engineered conventional phosphomimetic mutations at these sites. These mutations exhibited phenotypes associated with loss of CDC-20-activated APC/C function, including compromised osmotic integrity, that precluded analysis of one-cell stage embryos (Kim et al., 2017). We therefore took a different approach to generate more persistently phosphorylated CDC-20 based on the fact that threonine is conserved as the critical CDC-20 target site (**Fig. 1A**), combined with evidence that protein phosphatases prefer threonine over serine as a substrate (Cundell et al., 2016; Hein et al., 2017). To make CDC-20 more persistently phosphorylated, we therefore mutated Thr32 of CDC-20 to the less phosphatase-preferred serine. This subtle alteration in CDC-20 significantly increased mitotic duration, with NEBD to anaphase onset taking on average ~360s, compared to ~200s in wild-type embryos (**Fig. 1B-D**), indicating delayed APC/C activation. Notably, the effects of the two CDC-20 mutants on mitotic duration were independent of the spindle checkpoint, as the same effects were observed in the absence of MAD-3, a critical checkpoint effector (Lara-Gonzalez, Pines, et al., 2021; Musacchio, 2015)(**Fig. 1E**). These results establish that phosphoregulation of CDC-20 is a spindle checkpoint-independent mechanism controlling mitotic duration during embryonic divisions.

**Figure 1.**
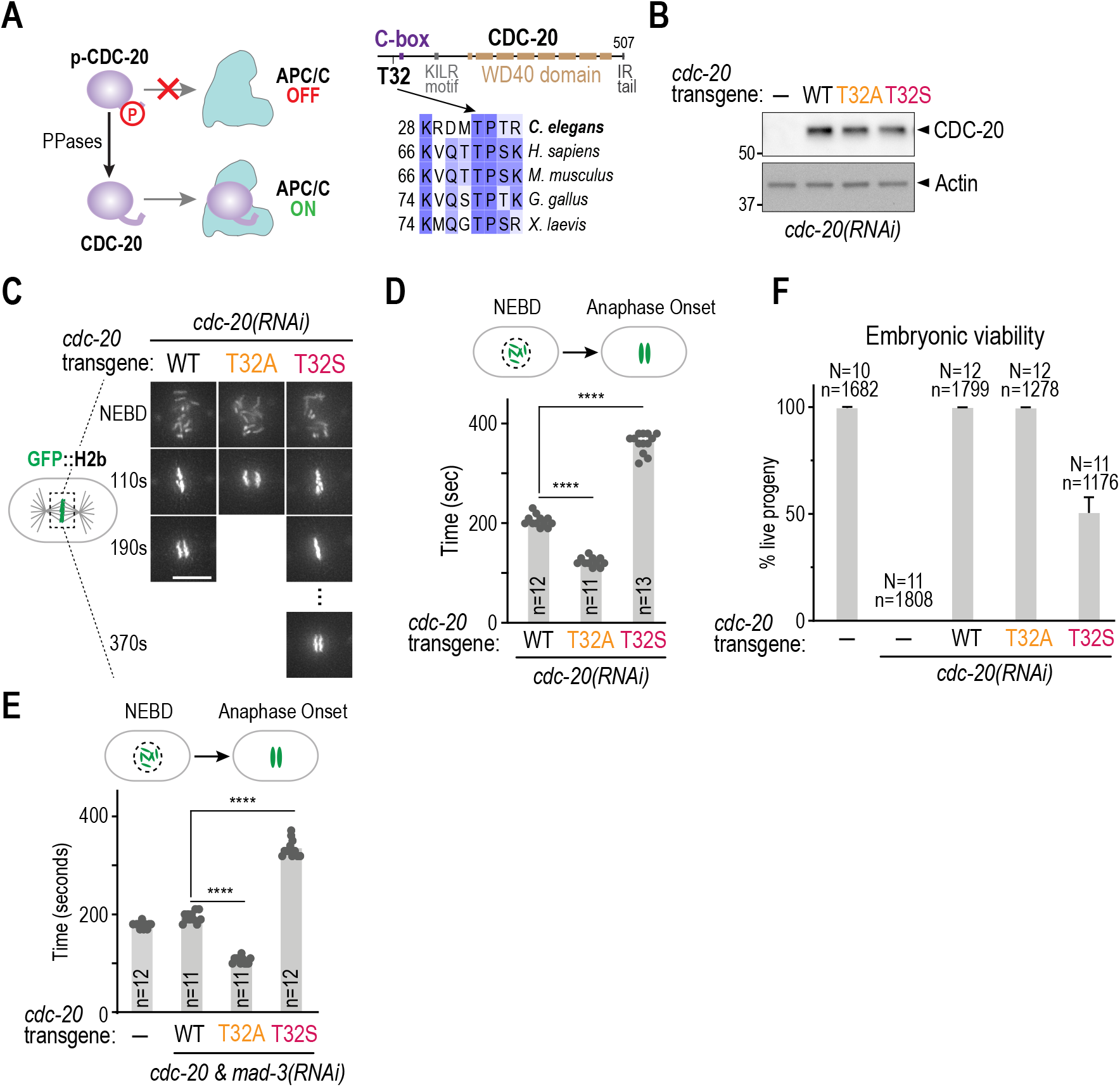
CDC-20 phosphoregulation controls early embryonic mitotic duration independently of the spindle checkpoint. **(A)** (*left*) Schematic of CDC-20 phosphoregulation based on biochemical analysis (Kim et al., 2017); phosphorylation reduces CDC-20 binding to APC/C. (*right*) Sequence alignments highlighting the conserved Thr32 Cdk phosphosite. **(B)** Immunoblot of lysates from worms with the indicated RNAi-resistant *cdc-20* mutant transgenes after depletion of endogenous CDC-20; actin serves as a loading control. **(C)** Stills from timelapse movies imaging chromosomes in embryos from the indicated conditions. Scale bar is 5 μm. **(D) & (E)** Quantification of the NEBD-anaphase onset interval for the indicated conditions. Bar height is the mean value. *n* is the number of embryos quantified. **(F)** Embryonic viability for the indicated conditions. *N* is number of worms whose progeny were scored and n is the number of embryos; mean and 95% confidence interval are plotted. p-values are from unpaired t-tests; ****: p<0.0001.

While acceleration of mitosis observed in Thr32Ala CDC-20 did not result in lethality (**Fig. 1F**), it exhibited synthetic larval lethality with spindle checkpoint inhibition (Kim et al., 2017). By contrast, extension of mitotic duration in Thr32Ser CDC-20 was by itself deleterious for embryonic development, as indicated by the significant embryonic lethality (**Fig. 1F**). Neither the meiotic arrest and osmotic integrity defects caused by loss of APC/C function (Golden et al., 2000; Rappleye et al., 2002), nor chromosome missegregation was observed in CDC-20 Thr32 mutants (*not shown*). These results suggest that the mitotic duration observed in wild-type embryos, which is defined by dynamic changes in CDC-20 phosphorylation, represents an optimum needed to support robust development.

### PLK-1 docking on BUB-1 promotes CDC-20 kinetochore localization and timely mitotic progression

CDC-20, but not the core APC/C, rapidly fluxes through kinetochores (Kallio et al., 2002; Kim et al., 2017). In prior work we showed that preventing CDC-20 from localizing to kinetochores or preventing CDC-20 dephosphorylation by removing the major kinetochore-localized phosphatase in *C. elegans* (PP1 bound to KNL-1), both extended the NEBD to anaphase onset interval to ~300s (Espeut et al., 2012; Kim et al., 2017). Combining these perturbations with mutation of CDC-20 N-terminal Cdk sites to alanines suppressed the extension of mitotic duration, indicating that CDC-20 recruitment to kinetochores promotes its dephosphorylation by PP1 (Kim et al., 2017). Preventing CDC-20 kinetochore recruitment extended mitosis to a lesser degree than what is observed with CDC-20 Thr32Ser and resulted in embryos that were viable but sensitized in their development to mild APC/C inhibition (Kim et al., 2017). These data suggest that localized CDC-20 dephosphorylation by kinetochore-recruited PP1 contributes to timely anaphase onset but that CDC-20 can also be dephosphorylated in the cytosol at a slower rate, with both mechanisms likely contributing to the mitotic timing optimum that is needed to support robust embryonic development.

The key binding site for CDC-20 at the kinetochore is the conserved ABBA motif in BUB-1, which interfaces with the structured WD40 domain of CDC-20 (**Fig. 2A**) (Di Fiore et al., 2015; Diaz-Martinez et al., 2015; He et al., 2013; Kim et al., 2017). In the process of analyzing other conserved regions of BUB-1, we discovered that a conserved “STP” motif located ~90 aa C-terminal to the ABBA motif, where Cdk phosphorylation creates a binding site for the Polo box domain of PLK-1 (Nguyen et al., 2021; Qi et al., 2006; Singh et al., 2021), is also required for CDC-20 to localize to kinetochores (**Fig. 2A**; BUB-1 PD^mut^, for PLK-1–docking mutant). Using purified fragments of BUB-1 and PLK-1, the Polo-docking motif in *C. elegans* BUB-1 has been validated as interacting with the Polo box domain in a Thr527 phosphorylation-dependent manner (SJP Taylor & F Pelisch, *personal communication*). PD^mut^ BUB-1 increased the NEBD to anaphase onset interval to exactly the same extent as selective mutation of the BUB-1 ABBA motif (**Fig. 2B**). PD^mut^ BUB-1 reduced but did not eliminate PLK-1 kinetochore localization, indicating the presence of an additional kinetochore-localized pool of PLK-1 (**Fig. 2C**). PD^mut^ BUB-1::GFP, which was expressed similarly to WT BUB-1::GFP, localized to kinetochores with a maximal level that was ~50% of WT BUB-1::GFP (**Fig. 2D; Fig. S1A**); this partial reduction is potentially due to reduced phosphorylation of KNL-1, the phosphoregulated kinetochore receptor of BUB-1 (Espeut et al., 2015; Moyle et al., 2014). Nevertheless, the reduction of kinetochore-localized PD^mut^ BUB-1 is unlikely to account for the NEBD-anaphase onset delay, as prior work reducing BUB-1 levels to a similar extent through deletion of binding sites in KNL-1 only mildly increased mitotic duration (Kim et al., 2015; Moyle et al., 2014). We therefore directly analyzed CDC-20 kinetochore localization and found that it was undetectable in PD^mut^ BUB-1 (**Fig. 2E**). Thus, the PLK-1 docking site on BUB-1 plays an essential role in recruiting CDC-20 to kinetochores. Consistent with this, similarly to ABBA^mut^ BUB-1, PD^mut^ BUB-1 was deficient for spindle checkpoint signaling (**Fig. S1B**), which requires CDC-20 recruitment to unattached kinetochores, where its association with MAD-2 is catalyzed to drive formation of mitotic checkpoint complexes (Lara-Gonzalez, Kim, et al., 2021; Piano et al., 2021). These data indicate that the PLK-1 docking site on BUB-1, while contributing to kinetochore pools of PLK-1 and BUB-1, is essential for CDC-20 kinetochore recruitment.

**Figure 2.**
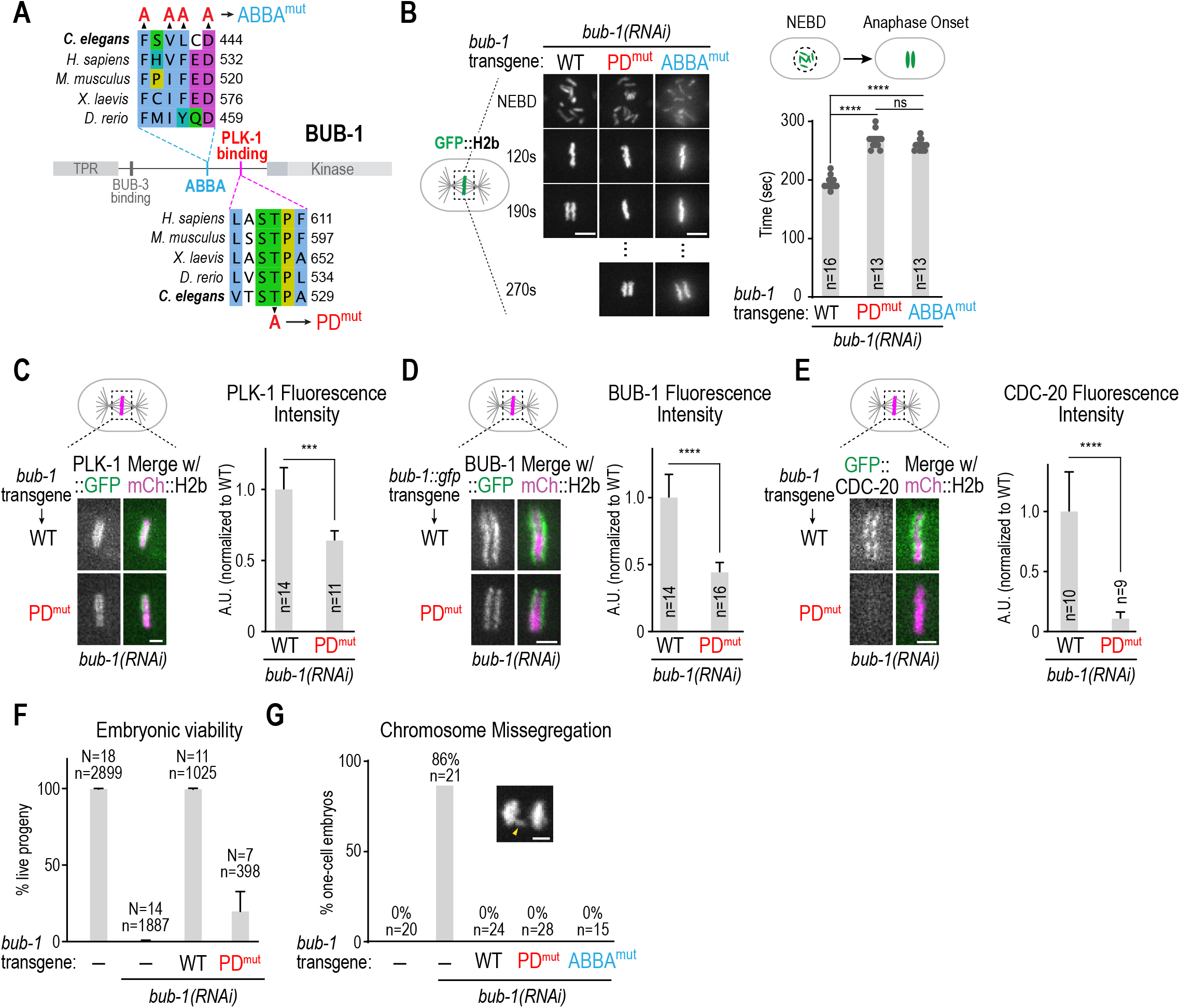
The PLK-1-docking site in BUB-1 is required for timely mitosis and CDC-20 kinetochore recruitment. **(A)** Sequence alignments of the CDC-20 binding ABBA motif and the PLK-1 docking motif in BUB-1. Mutations engineered to disrupt CDC-20 binding (ABBA^mut^) and PLK-1 interaction (PD^mut^) are indicated. **(B)** (*left*) Images from timelapse movies of chromosomes for the indicated conditions. Images of WT and PD^mut^ BUB-1 were taken using an Andor Revolution confocal system; images of ABBA^mut^ BUB-1 were taken using a Deltavision widefield deconvolution microscope. (*right*) NEBD-anaphase onset interval for the indicated conditions. **(C-E)** Representative images (*left*) and quantification of chromosomal fluorescence intensity at metaphase (*right*) in WT vs PD^mut^ BUB-1 for the indicated proteins. *n* is the number of embryos quantified. Error bars are the 95% confidence interval. Scale bar is 2 μm. **(F)** Embryonic viability analysis for the indicated conditions. *N* is number of worms analyzed; *n* is total number of progeny scored. Error bars are the 95% confidence interval. **(G)** Frequency of anaphase chromosome missegregation scored from timelapse movies for the indicated conditions. Inset shows an example of lagging anaphase chromatin (*arrowhead*). *n* is the number of embryos quantified. Scale bar is 2 μm. p-values are from unpaired t-tests; ****: p<0.0001.

ABBA^mut^ BUB-1 on its own does not exhibit significant embryonic lethality but is sensitive to mild APC/C inhibition (Kim et al., 2017). By contrast, PD^mut^ BUB-1 exhibited significant embryonic lethality on its own (**Fig. 2F**). BUB-1 inhibition leads to penetrant embryonic lethality and visible chromosome missegregation characterized by anaphase bridges (Essex et al., 2009; Kim et al., 2015). Anaphase chromosome missegregation was not observed for either PD^mut^ or ABBA^mut^ in one-cell embryos (**Fig. 2G**), suggesting that chromosome missegregation was unlikely to be cause of the lethality observed with PD^mut^ BUB-1. Whether the lethality observed in PD^mut^ BUB-1 is due to impacts on mitosis or because it reflects an as-yet-unknown role for BUB-1-bound PLK-1 during embryonic development, awaits future analysis.

### PLK-1 kinase activity is required for CDC-20 kinetochore localization

The analysis of PD^mut^ BUB-1 suggested that PLK-1 docked onto BUB-1 is important to recruit CDC-20 to kinetochores. We next sought to address if PLK-1 kinase activity is important for this function. As PLK-1 is required for meiotic progression and mitotic entry (Chase et al., 2000), mutants that eliminate its kinase activity cannot be analyzed. We therefore used an analog-sensitive (AS) allele of PLK-1 engineered at the endogenous locus using genome-editing (Gomez-Cavazos et al., 2020). To temporally control activity inhibition, we employed a method to permeabilize embryos (Carvalho et al., 2011) and added the inhibitor specifically targeting analog-sensitive PLK-1 to permeabilized embryos immobilized in microwells around the time of NEBD (**Fig. 3A**). We conducted this analysis in strains expressing *in situ* tagged BUB-1::GFP or transgene-encoded GFP::CDC-20. When PLK-1 was wild-type, inhibitor addition had no effect on either BUB-1 or CDC-20 kinetochore localization (**Fig. 3B,C**). By contrast, following inhibitor addition in PLK-1^AS^, BUB-1 was still kinetochore-localized but CDC-20 kinetochore localization was significantly diminished (**Fig. 3B,C; Fig. S1C**). Thus, PLK-1 kinase activity is important for CDC-20 kinetochore recruitment.

**Figure 3.**
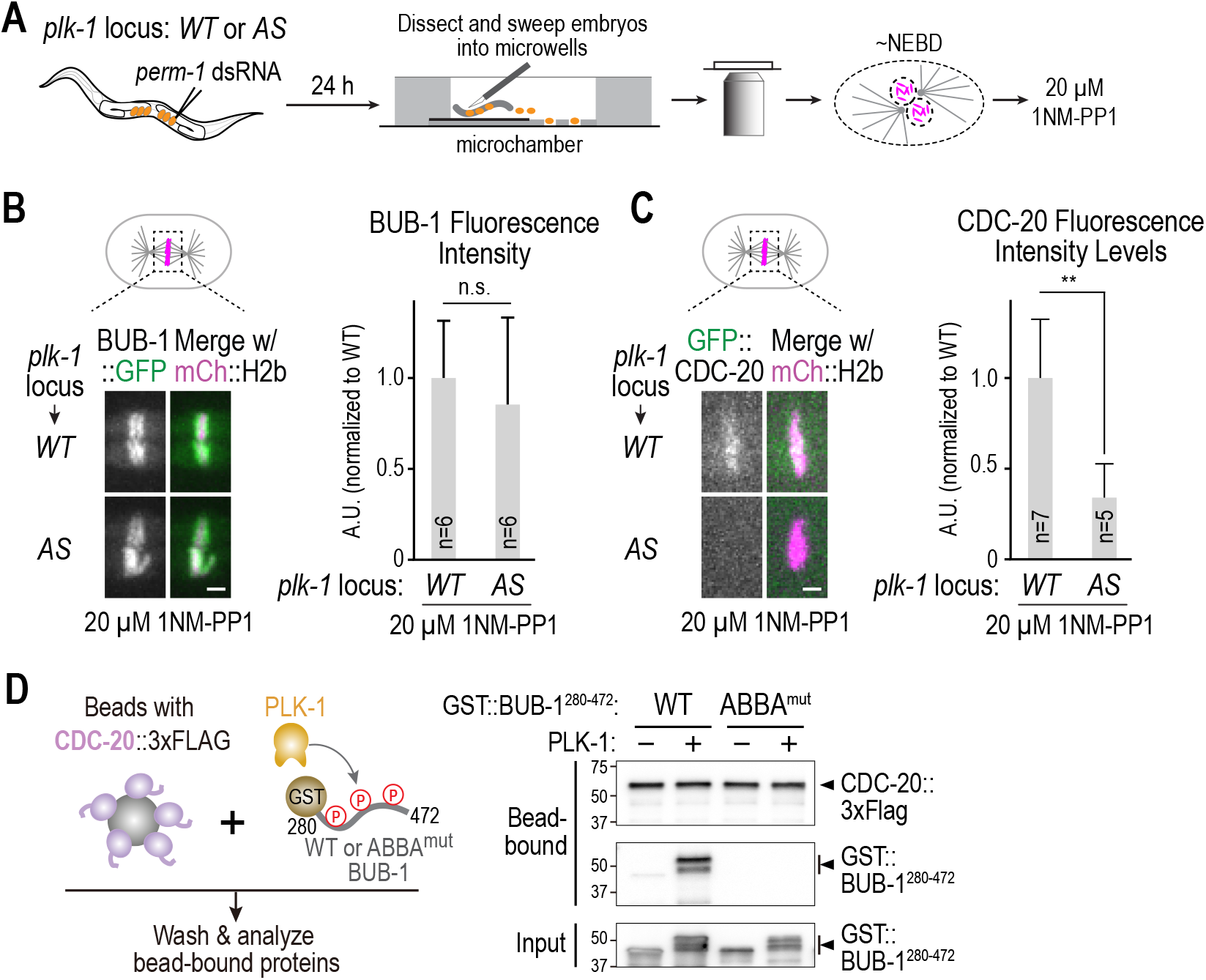
PLK-1 kinase activity is important for CDC-20 kinetochore recruitment. **(A)** Schematic of assay used to permeabilize embryos and temporally control PLK-1 inhibition using an analog-sensitive (AS) allele engineered at the endogenous *plk-1* locus. **(B & C)** Representative images (*left*) and quantification of chromosomal fluorescence intensity (*right*) for the indicated conditions. *n* is the number of embryos quantified. Scale bars, 2 μm. p-values are from t-tests; n.s.: not significant; **: p<0.01. **(D)** (*left*) Schematic of assay used to assess interaction of an ABBA motif-containing BUB-1 fragment with full-length CDC-20; constitutively-active purified PLK-1 was used phosphorylate the BUB-1 fragment. (*right*) Immunoblot of bead-bound proteins (*top*) and PLK-1 phosphorylated BUB-1 fragment input into the binding reaction (*bottom*). Results shown are representative of two independent experiments.

### PLK-1 phosphorylation promotes BUB-1’s interaction with CDC-20

The analysis of PD^mut^ BUB-1 and PLK-1 kinase activity inhibition suggests that the kinase activity of PLK-1 tethered on BUB-1 is essential to recruit CDC-20 to kinetochores. As the primary binding site for CDC-20 at kinetochores is BUB-1’s ABBA motif, which is ~90 aa N-terminal to the PLK-1 docking site, an attractive model is that tethered PLK-1 phosphorylates residues in BUB-1 to enhance the ABBA-dependent interaction with CDC-20. To test this model, we expressed full-length *C. elegans* CDC-20 in a human cell protein expression system, bound it to beads, and combined the CDC-20-coated beads with a GST-fused BUB-1 fragment (aa 280-472) that contains the ABBA motif. Without PLK-1, the ABBA-containing BUB-1 fragment interacted weakly with CDC-20. By contrast, incubation of the BUB-1 fragment with PLK-1 (constitutively active T194D *C. elegans* PLK-1) in the presence of ATP strongly stimulated its binding to CDC-20 (**Fig. 3D**). Robust PLK-1 phosphorylation was evident in the retarded gel mobility of the GST-BUB-1^280-472^ fragment (**Fig. 3D**). Importantly, mutating the critical residues in the ABBA motif that are essential for interaction with the WD40 domain of CDC-20 eliminated the PLK-1 phosphorylation-stimulated interaction, even though significant phosphorylation of the ABBA^mut^ BUB-1 fragment was observed (**Fig. 3D**). These biochemical data indicate that PLK-1 phosphorylation directly promotes the ABBA-dependent interaction of CDC-20 with BUB-1.

### An unbiased screen identifies putative target sites of PLK-1 in BUB-1 that contribute to CDC-20 kinetochore localization

In parallel to developing the biochemical assay monitoring PLK-1 stimulated BUB-1–CDC-20 interaction, we pursued an *in vivo* approach to screen for key target sites of PLK-1 kinase activity in BUB-1. Based on the ability to replace endogenous BUB-1 with precisely engineered mutants, we employed an unbiased mutagenesis strategy that previously yielded PLK-1 target sites important for cytokinesis and spindle assembly (Gomez-Cavazos et al., 2020; Ohta et al., 2021). Specifically, we defined 45 putative PLK-1 sites in BUB-1 using phosphorylation consensus motif and conservation criteria (**Fig. S2A**). We mutated these 45 sites in 8 clusters (**Fig. 4A; Fig. S2B**); RNAi-resistant untagged BUB-1 transgenes encoding the 8 cluster mutants were integrated in single-copy and crossed into a strain expressing GFP::CDC-20. Following endogenous BUB-1 depletion, CDC-20 localization was analyzed by live imaging. This screening approach revealed that the Cluster V mutant, in which 7 putative PLK-1 sites were mutated to alanine, has significantly reduced CDC-20 kinetochore localization (**Fig. 4B,C**). One possible explanation for reduced CDC-20 localization is that the kinetochore localization of the Cluster V mutant BUB-1 was itself reduced. As transgene-encoded BUB-1 was untagged, in order to exclude this possibility we monitored localization of GFP::BUB-3, which is dependent on BUB-1 for its kinetochore localization (Essex et al., 2009; Kim et al., 2015). BUB-3 localization in the Cluster V mutant was indistinguishable from controls (**Fig. S2C**); in addition, the Cluster V mutant, all well as all other cluster mutants, rescued the embryonic lethality caused by BUB-1 depletion (**Fig. S2D**). These results indicate that one or more target sites in Cluster V are important for CDC-20 kinetochore localization.

**Figure 4.**
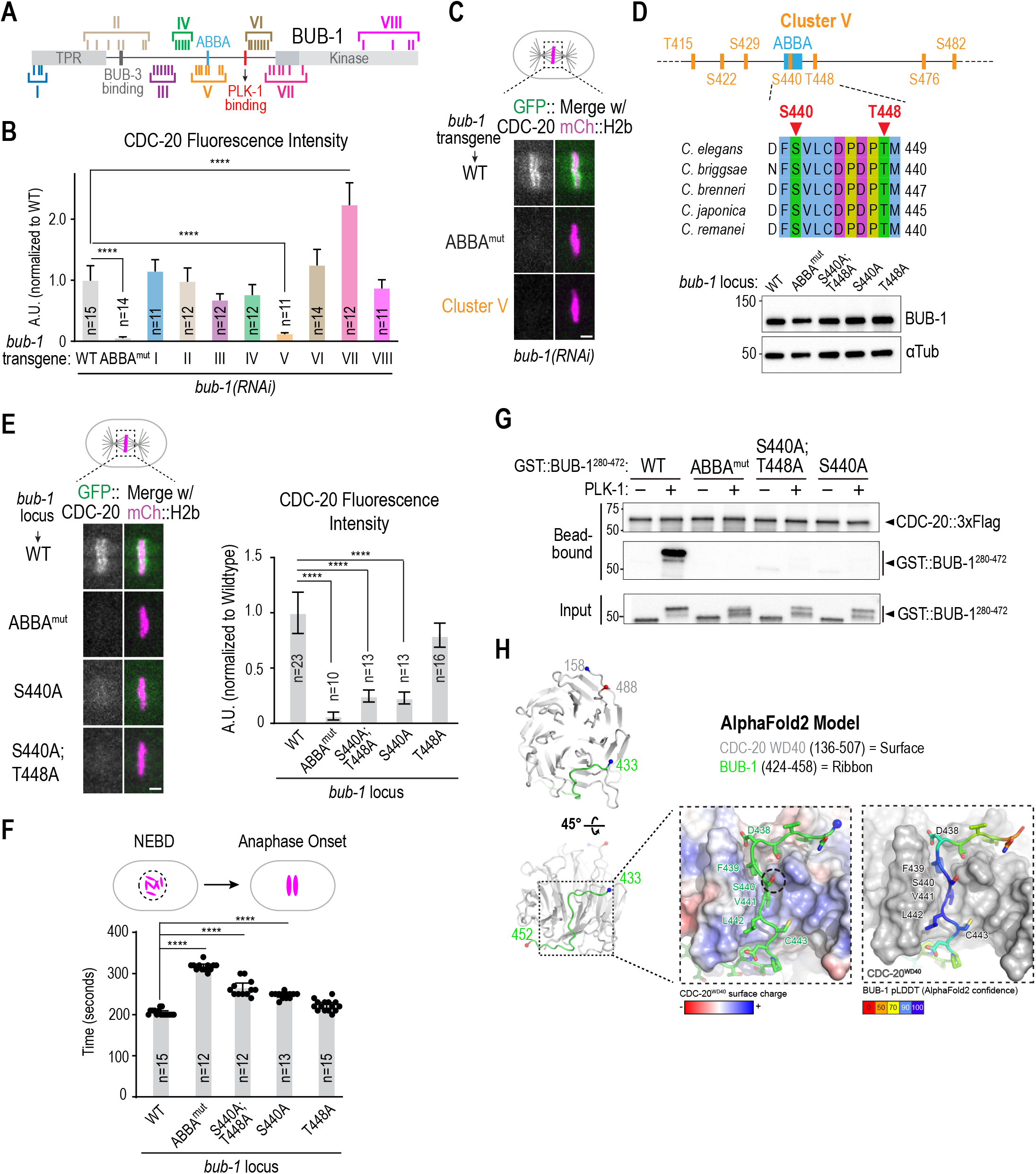
Mechanism by which PLK-1 phosphorylation enhances affinity of the BUB-1 ABBA motif for CDC-20 to ensure timely mitosis. **(A)** Schematic of BUB-1 highlighting 45 putative PLK-1 target sites and 8 clustered alanine mutants designed to assess effects on CDC-20 kinetochore recruitment. **(B)** Quantification of GFP::CDC-20 localization for the indicated conditions. *n* is the number of embryos quantified. p-values are from unpaired t-tests; ****: p<0.0001. **(C)** Representative images of CDC-20 localization in the indicated conditions. Scale bar, 2 μm. **(D)** (*top*) Details on sites mutated in Cluster V, which is centered around the ABBA motif. (*bottom*) Immunoblot of BUB-1 following genome-editing of the endogenous *bub-1* locus to introduce the indicated mutations. The genome editing was conducted in a strain with transgene-encoded GFP::CDC-20 and mCherry::H2b. **(E)** Analysis of CDC-20 localization in the indicated conditions. *n* is the number of embryos quantified. Scale bar, 2 μm. p-values are from unpaired t-tests; ****: p<0.0001. **(F)** Quantification of NEBD-anaphase onset interval for the indicated conditions. *n* is the number of embryos quantified. Error bars are the 95% confidence interval. p-values are from unpaired t-tests; ****: p<0.0001. **(G)** Binding assay conducted as in *Fig. 3D*, except that specific phospho-site mutant GST::BUB-1 fragments were employed, in addition to WT and ABBA^mut^. Immunoblots on top shows protein bound to CDC-20-coated beads; bottom immunoblot is of the GST::BUB-1 fragments phosphorylated by PLK-1. Results shown are representative of two independent experiments..**(H)** AlphaFold2 model of *C. elegans* BUB-1 ABBA motif and *C. elegans* CDC-20. Two views show the overall position of the BUB-1 ABBA motif (green) docked against the WD40 beta-propeller domain (grey) in CDC-20. The start and ends of the shown chains are represented with blue and red colored spheres, respectively. The enlarged, space-filled zoom views show: overall surface charge on ABBA binding site on CDC-20 WD40 (*left*); confidence score of BUB-1 ABBA motif residues in this binding site (*right*). The Ser440 residue is outlined by a black dotted circle.

Interestingly, the Cluster VII mutant, which alters a number of residues in the N-terminal extension involved in Bub1 kinase activation (Kang et al., 2008), was associated with elevated CDC-20 kinetochore localization (**Fig. 4B**; **Fig. S2E**), an unexpected result that will be investigated further in future work. Collectively, *in vivo* screening of putative PLK-1 sites in BUB-1 identified Cluster V, which is centered around the ABBA motif, as being important for CDC-20 kinetochore localization. This finding is consistent with the biochemical data showing that PLK-1 phosphorylation of a fragment of BUB-1 containing the ABBA motif is sufficient to stimulate direction interaction with CDC-20 (**Fig. 3E**).

### A PLK-1 target site in the CDC-20 binding motif of BUB-1 makes a major contribution to CDC-20 interaction, CDC-20 kinetochore localization, and timely mitotic progression

We next focused on which of the 7 mutated PLK-1 sites in Cluster V are critical for CDC-20 kinetochore recruitment. Since sequence alignments indicated that Ser440 and Thr448 were conserved across *Caenorhabditis* species (**Fig. 4D**), we mutated them to alanine, individually and in combination, at the endogenous *bub-1* locus in a strain expressing GFP::CDC-20 and mCherry::H2b; all 3 homozygous mutants were viable and had normal BUB-1 levels (**Fig. 4D**). Further analysis indicated that Ser440, which is within the ABBA motif, makes a major contribution to CDC-20 kinetochore recruitment, whereas mutation of Thr448 has a significantly milder effect (**Fig. 4E**). Of the other sites, Ser422, Ser476 and Ser482 were not conserved across *Caenorhabditis* species, and while Thr415 was conserved, mutation of the threonine to alanine had no effect on CDC-20 kinetochore localization (*not shown*). Notably neither the Ser440Ala nor the Ser440Ala/Thr488Ala mutants of BUB-1 reduced CDC-20 kinetochore localization to the same extent as the ABBA mutant (**Fig. 4E**). This observation is consistent with the notion that the ABBA motif of BUB-1 has intrinsic affinity for CDC-20 that is enhanced by local PLK-1 phosphorylation.

The strong effect of mutation of Ser440 of BUB-1 on CDC-20 kinetochore localization predicts that mitotic progression will be delayed in this mutant. Analysis of mitotic timing revealed that the Ser440Ala mutant significantly increased the NEBD-Anaphase onset interval but the magnitude of the increase was less than for ABBA^mut^ or PD^mut^ BUB-1 (**Fig. 4F**). The Thr448Ala mutant on its own did not significantly extend mitotic duration but the Ser440Ala/Thr488Ala mutant was slightly elevated relative to Ser440Ala mutant alone (**Fig. 4F**). We note that PD^mut^ BUB-1 significantly reduced levels of BUB-1 and PLK-1 at kinetochores and that, in contrast to ABBA^mut^, the Ser440Ala and the Ser440Ala/Thr488Ala mutant reduce but do not eliminate CDC-20 kinetochore localization (**Fig 4B)**. These observations are consistent with the model that Ser440 is a major target site of PLK-1 whose phosphorylation enhances the intrinsic affinity of the BUB-1 ABBA motif for CDC-20.

The data above predict that Ser440 should be important for the PLK-1 phosphorylation-stimulated interaction between purified BUB-1 and CDC-20. CDC-20 interaction analysis following PLK-1 phosphorylation of the GST::BUB-1 fragment containing the ABBA motif after mutating Ser440 alone or both Ser440 and Thr448 revealed that preventing Ser440 phosphorylation greatly reduced the PLK-1 stimulated interaction between BUB-1 and CDC-20 (**Fig. 4G**). Analysis of the input indicated that the Ser440Ala and the Ser440Ala/Thr488Ala mutant BUB-1 fragments were phosphorylated by PLK-1, as evidenced by their retarded gel mobility, highlighting the specific importance of Ser440 in the PLK-1 activity-stimulated interaction (**Fig. 4G**). The difference in mobility between the mutant and WT GST::BUB-1 fragments is also suggestive of phosphorylation of the mutated sites by PLK-1. Mass spectrometry of PLK-1 phosphorylated WT BUB-1 fragment confirmed phosphorylation of Thr448; unfortunately, despite multiple attempts with different protocols, the ABBA motif peptide including Ser440 was never detected (*not shown*). Nonetheless, the balance of *in vitro* and *in vivo* evidence strongly supports the conclusion that phosphorylation of Ser440 by PLK-1 significantly enhances the affinity of BUB-1 for CDC-20.

An Alphafold2 model of a *C. elegans* CDC-20-BUB–1 complex revealed that the BUB-1 ABBA motif residues interact with the β-propeller WD40 domain of CDC-20 in the inter-blade groove region (**Fig. 4H; Fig. S3A**;(Evans et al., 2022)); structural superposition indicated that the modeled binding of *C. elegans* BUB-1 ABBA motif to *C. elegans* CDC-20 is equivalent to the structurally characterized interaction of *S. cerevisiae* Acm1’s ABBA motif with the WD40 domain of the Cdc20 ortholog Cdh1 (**Fig. S3B**;(He et al., 2013)). Notably, Ser440 of BUB-1 is positioned in the vicinity of a basic region on *C. elegans* CDC-20 (specifically Arg234), potentially suggesting favorable interactions in its phosphorylated state.

## DISCUSSION

In summary, the results shown here suggest that a specific pool of PLK-1 tethered ~90 amino acids distal to the CDC-20 binding ABBA motif in BUB-1 phosphorylates this conserved motif to promote CDC-20 flux through kinetochores in *C. elegans* embryos. ABBA motifs are present in multiple Cdc20-interacting proteins, including cyclin A, the Bub1-related protein BubR1, and Acm1 (Burton et al., 2011; Di Fiore et al., 2015; Diaz-Martinez et al., 2015). While a Serine residue analogous to Ser440 is not present in vertebrate Bub1 ABBA motifs (**Fig. 2A)**, a Serine is present at this location in the ABBA motifs of BubR1 and cyclin A (Di Fiore et al., 2015). Whether phosphorylation of this Serine or in residues adjacent to the ABBA motif enhances the affinity of Cdc20-interacting proteins in other species, analogous to what we report here for BUB-1 in *C. elegans*, will be important to address in the future.

CDC-20 flux through kinetochores is important for both mitotic progression, via its dephosphorylation, and for delaying mitosis when kinetochores are persistently unattached; this latter function requires incorporation of CDC-20 into mitotic checkpoint complexes and is also compromised by loss of the BUB-1 tethered pool of PLK-1. Notably, CDC-20 phosphoregulation at the kinetochore and elsewhere functions as a means of controlling mitotic duration independent of the spindle checkpoint, the best-studied mechanism pacing mitosis, and may be particularly significant when mitosis must be executed in a rapid time frame.

## ACKNOWLEDGMENTS

We thank members of the Oegema and Desai labs for their support, Federico Pelisch for sharing unpublished work and helpful discussions, Jeffrey Woodruff for the gift of constitutively active PLK-1, and Aleesa Schlientz for comments on the manuscript. This work was supported by a grant from the NIH to A.D. (R01 GM074215), and a grant from the NIH to K.D.C. (R35 GM144121). J. H. was supported by an NSF Graduate Research Fellowship (grant #1650112). A.D. and K.O. acknowledge salary support from the Ludwig Institute for Cancer Research.

## AUTHOR CONTRIBUTIONS

T.K. and A.D. initiated the project. T.K. made the initial observations that were pursued by J.H., who conducted a majority of the experiments with significant guidance from P.L-G.; M.O. provided key expertise for the biochemical interaction analysis; J.S.G.-C. conducted the inhibitor treatment experiments; K.O. and J.S.C-G. helped define putative PLK-1 sites and design the cluster mutagenesis strategy; A.D. and K.D.C. conducted structural modeling and provided advice on mutation engineering; J.H. and P.L-G. prepared initial figure and text drafts; J.H., P.L-G., K.O., T.K.. and A.D. worked on finalizing the manuscript, with input from all authors.

## SUPPLEMENTAL FIGURE LEGENDS

**Figure S1.**
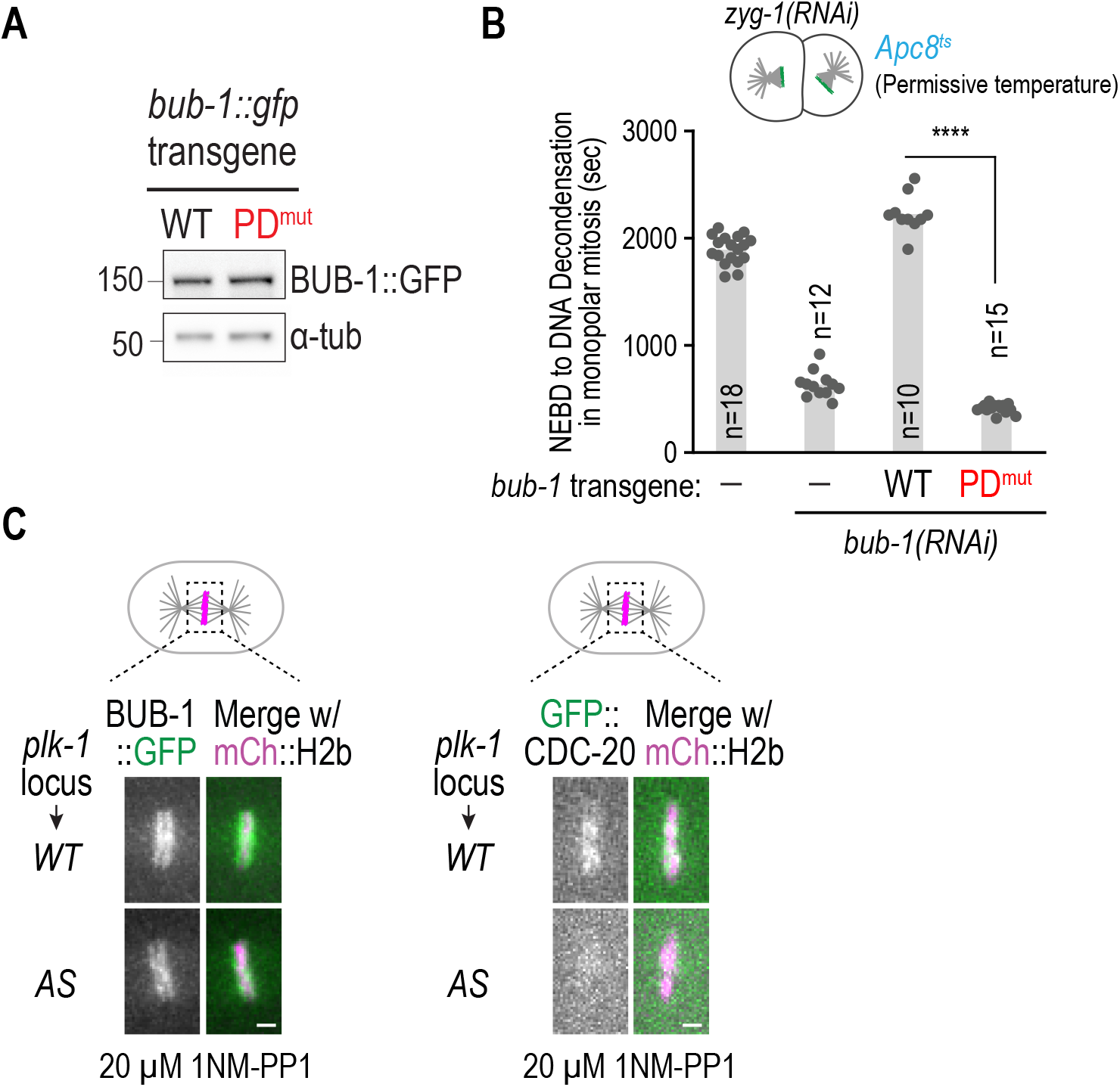
Additional analysis of PLK-1 docking mutant BUB-1 and of PLK-1 kinase activity inhibition. **(A)** Immunoblot of WT and PD^mut^ BUB-1::GFP expressed from single-copy transgene insertions. PD^mut^ BUB-1 is expressed at similar level to WT BUB-1. α-tubulin serves as a loading control. **(B)** Analysis of spindle checkpoint signaling in PD^mut^ BUB-1. Spindle checkpoint signaling at unattached kinetochores was analyzed using monopolar spindle formation in 2-cell embryos induced by depletion of ZYG-1, which is essential for centriole duplication; *Apc8^ts^* at the permissive temperature was used to enhance sensitivity of the checkpoint assay (Kim et al., 2017). Time from NEBD to chromosome decondensation was monitored from timelapse movies of GFP::H2b. Depletion of BUB-1 abolishes checkpoint signaling and is rescued by expression of RNAi-resistant transgene-encoded WT BUB-1. By contrast, PD^mut^ BUB-1 failed to rescue, which is consistent with the inability of CDC-20 to localize to kinetochores in this mutant. *n* is the number of embryos quantified. p-values are from unpaired t-tests; ****: p<0.0001. **(C)** Additional examples of BUB-1::GFP and GFP::CDC-20 localization following temporally-controlled PLK-1 inhibition. Scale bar, 2 μm.

**Figure S2.**
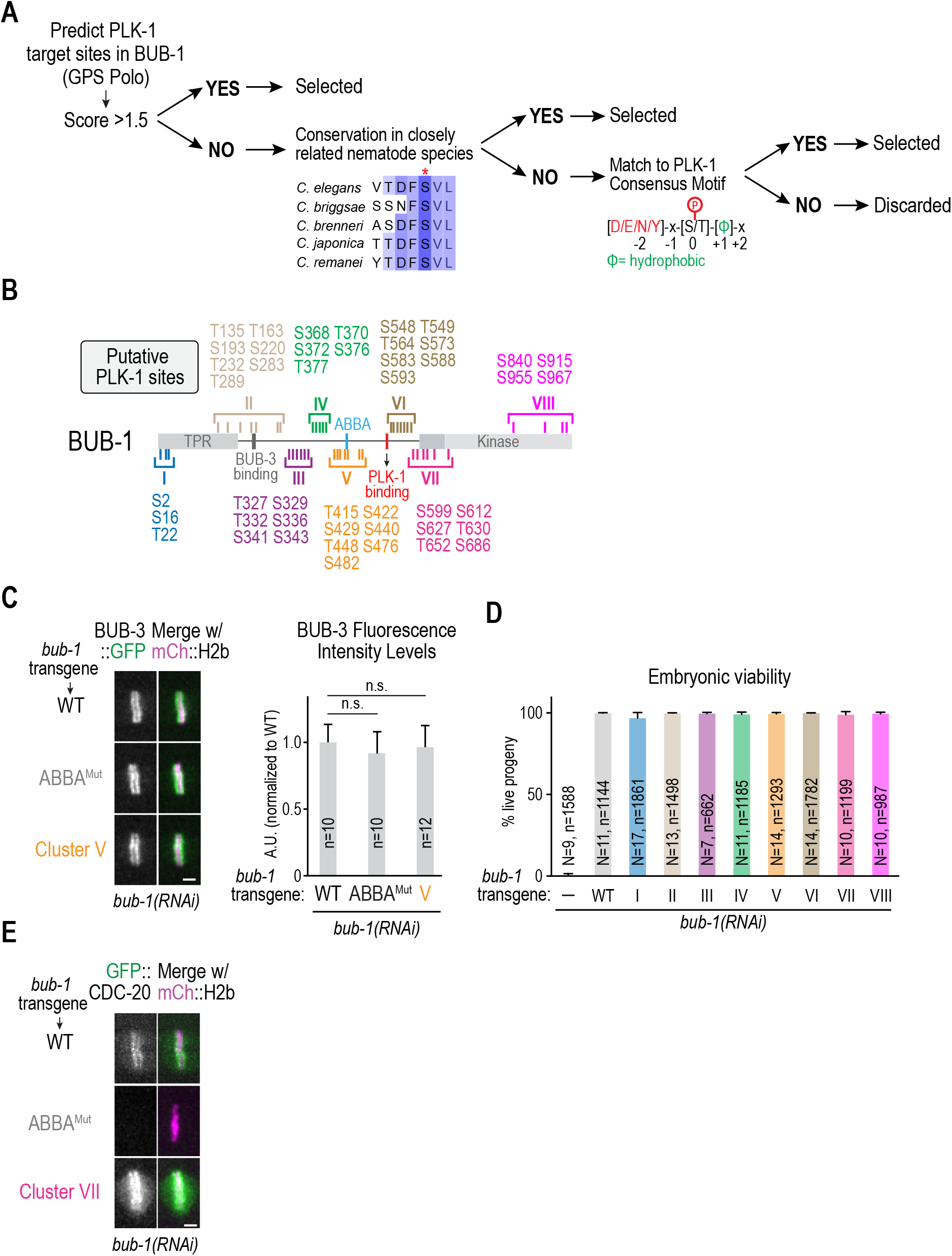
Selection and analysis of putative PLK-1 sites in BUB-1. **(A)** Schematic of approach used to select 45 putative PLK-1 target sites in BUB-1. **(B)** List of sites in the 8 clusters that were mutated to alanine. **(C)** Analysis of BUB-3::GFP localization in the indicated conditions. (*left*) Example images of BUB-3::GFP localization; (*right*) quantification of BUB-3::GFP signal on aligned chromosomes. BUB-3 requires BUB-1 for its kinetochore localization; normal BUB-3 localization in the Cluster V mutant indicates that the 7 sites mutated do not affect BUB-1 kinetochore localization. **(D)** Embryonic viability analysis for the indicated conditions. N is number of worms analyzed. *n* is the number of embryos quantified. Error bars are the 95% confidence interval. **(E)** Image of CDC-20 hyper-localization in the Cluster VII mutant. For quantification, see *Fig. 4B. n* is the number of embryos quantified. Scale bar, 2 μm. p-values are from unpaired t-tests; n.s.= not significant.

**Figure S3.**
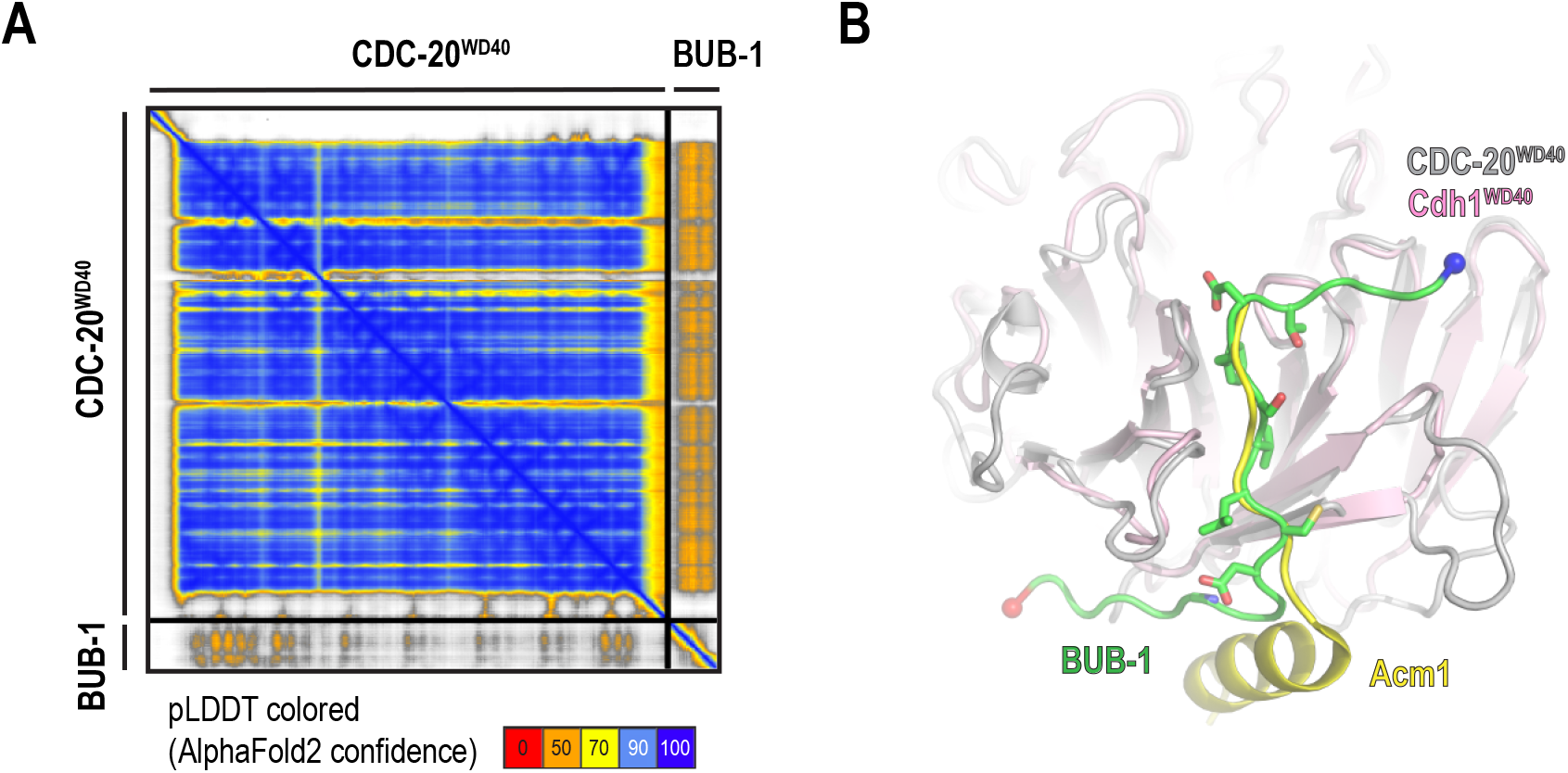
Alphafold model of CDC-20 WD40 domain interaction with the BUB-1 ABBA region. **(A)** Predicted alignment error (PAE) plot of the confidence of the AlphaFold2 model shown in *Fig. 4H*. **(B)** Structural superposition of *C. elegans* CDC-20 WD40 domain::BUB-1 ABBA motif model with the structure of *S. cerevisiae* Cdh1::Acm1 A motif (PDB ID 4BH6). The root mean square deviation for the superposition of S. cerevisiae Cdh1 WD40 with *C. elegans* CDC-20 WD40 is 0.76 Å for 212 Cα atoms.

## METHODS

### Worm Strains

*C. elegans* strains (**Table S1**) were maintained at 16-20°C. RNAi-resistant *bub-1* and *cdc-20* transgenes were previously described (Kim et al., 2017; Moyle et al., 2014). Transgenes were cloned into pCFJ151 or pCFJ352 and injected into strains EG6429 or EG6701, respectively, along with constructs encoding for the Mos transposase and a mix of four negative selection markers against extrachromosomal plasmid arrays. One week after injection, integrants were selected on the basis of their rescuing of the *unc* phenotype of the parental strains and their lack of negative selection markers, and further confirmed by PCR.

### RNAi (Table S2)

dsRNAs were synthesized *in vitro*. Briefly, target sequences were amplified by PCR using oligos containing either T3 or T7 promoters at their 5’ end. Complementary RNAs were synthesized from the PCR templates using T3 or T7 RNA polymerases (MEGAscript T3 or T7 kit, ThermoFisher). Single-stranded RNA products were then purified using MEGAclear kit (ThermoFisher) and annealed by incubation at 68°C for 10 minutes followed by 37°C for 30 minutes. Resulting dsRNA were measured by A_260nm_ using a nano-drop spectrophotometer.

For RNAi based depletions, L4 stage worms were injected with 0.8-1 mg/mL of dsRNA and incubated at 20°C for 36-48 hours before imaging-based experiments. For embryonic viability assays, L4 stage worms were injected with dsRNA and recovered at 20°C for 24 hours. Injected worms were singled and allowed to lay progeny for 24 hours at 20°C. Parental worms were then removed and after further 24 hr, progeny was scored as either viable or dead.

### CRISPR-Cas9 (Table S3)

CRISPR-Cas9 Ribonucleoproteins (RNPs) were used for *in situ* mutagenesis (Paix et al., 2017). Target sequences were identified using CHOPCHOP (https://chopchop.cbu.uib.no/). Purified Cas9 protein was purchased from MacroLab. Repair oligos were ordered from Eton Biosciences as HPLC-purified products. TracrRNA and crRNAs were ordered from IDT.

A co-CRISPR strategy was used to facilitate the identification of edits. For this, a guide RNA mix was created using 100 μM tracrRNA, 25 μM *dpy-10* crRNA, and 75 μM targeting crRNA. RNAs were annealed by incubation at 95 °C for 5 minutes, followed by hybridization at 22 °C for 5 minutes. Then, 2 μL of the RNA mix were mixed with 5 μL of 40 μM Cas9 protein and incubated at 22 °C for 5 minutes to generate RNP complexes. A repair oligo was added to the mix to a final concentration of 5.4 μM and the resulting mix was injected in the germline of adult worms. After 4-5 days, between 24-30 F1 progeny from plates with high proportion of *dpy-10* mutant worms (“jackpot” broods) were singled out, and edits were screened through PCR-based amplification of the allele, followed by sanger sequencing. Edited strains were outcrossed 1 time to the parental strain to remove the *dpy-10* mutated allele.

### Fluorescence imaging of *C. elegans* embryos

Gravid adult hermaphrodite worms were dissected in M9 buffer, early embryos were transferred onto a 2% agarose pad using a mouth pipet, covered with a 22×22 mm coverslip, and imaged at 20°C. For kinetochore localization assays, embryos were imaged on an Andor Revolution confocal system (Andor) coupled to a CSU-10 spinning disk confocal scanner (Yokogawa) and an electron multiplication back-thinned charge-coupled device camera (iXon, Andor), using either a 60x or 100x 1.4 NA Plan Apochromat objective. A 6×2 μm z-stack was collected every 10 or 20 seconds.

For mitotic timing experiments, timelapse imaging of one cell embryos expressing GFP::H2b or mCherry::H2b was performed on a widefield deconvolution microscope (DeltaVision) connected to a charge-coupled device camera (pco.edge 5.5 sCMOS; PCO) and a 60x 1.42NA PlanApo N objective (Olympus). A 5×2 μm z-stack was collected every 10 or 20 seconds.

### PLK-1 inhibition assays

The PLK-1 analog-sensitive (AS) allele (C52V and L115G) was previously described (Gomez-Cavazos et al., 2020). For PLK-1 inhibition experiments, L4 stage worms were injected with *perm-1* dsRNA to prevent formation of the drug-impermeable eggshell and incubated at 20 °C for 16-18 hours before imaging. Worms were dissected in custom made imaging chamber slides and embryos deposited into built-in wells. Embryos were monitored by DIC imaging, and 1NM-PP1 was added to the well to a final concentration of 20 μM after NEBD. Embryos were imaged every 20 seconds using 8×2 μm z stacks using DIC, RFP, and GFP channels. Embryo permeability was assessed at each imaging session by addition of lipophilic dye FM4-64, which binds to the plasma membrane of permeabilized but not impermeable embryos (Gomez-Cavazos et al., 2020).

### Protein Expression and Purification (Protein List Table S4)

GST tagged BUB-1 (280-472) wildtype and mutant constructs were cloned into pGEX-6P-1 vectors and transformed into Bl21(DE3)pLys *E. coli* cells. Cultures were grown at an OD_600_ of 0.4-0.6 and induced overnight at 20°C with 0.4 mM IPTG. The next morning, cells were collected by centrifugation and re-suspended with lysis buffer (25 mM HEPES pH 7.5, 300 mM NaCl, 1 mM MgCl_2_, 0.1% Triton X-100, 10 mM βME), supplemented with 1 mM PMSF, cOmplete EDTA protease inhibitors (Roche) and 1 mg/mL Lysozyme. Cells were lysed by sonication, and lysates were cleared by ultracentrifugation for 30 minutes at 40,000 rpm at 4°C in a 45Ti rotor. GST-tagged proteins were purified from the cleared lysates using glutathione-sepharose 4B beads (Cytiva), and incubated with end-over-end rotation for 2 hours at 4°C. Beads were washed 3 times with lysis buffer and eluted with 10 mM reduced glutathione in 50 mM Tris-HCl pH 8.0; 150 mM NaCl. Fractions containing the most protein were combined and dialyzed against storage buffer (25 mM HEPES pH 7.5, 200 mM NaCl, 1 mM MgCl_2_, 0.1% Triton X-100, 10 mM βME, 5% glycerol). Proteins were aliquoted, snap frozen in liquid nitrogen and stored at −80°C.

### *In vitro* binding assays

A construct encoding for CDC-20::3xFLAG under a CMV promoter was transfected into Freestyle 293-F cells (Thermo Fisher Scientific). 48 hours later, cells were harvested and washed with PBS before re-suspending in lysis buffer (20 mM Tris-Cl pH 7.5, 50 mM NaCl, 5 mM EGTA, 2 mM MgCl_2_, 0.5% Triton X-100, 1 mM DTT), supplemented with 1x cOmplete Protease inhibitor cocktail (Millipore). Cells were sonicated for 6 minutes in an ice-cold water bath, then centrifuged for 15 minutes at 15,000xg at 4°C. Whole-cell lysates were then incubated with Anti-FLAG beads (Sigma) for 2 hours at 4°C and beads were washed four times with lysis buffer.

Meanwhile, 2 μM GST::BUB-1 (280-472) was phosphorylated with 500 nM constitutively active PLK-1 T194D (gift from Jeffrey Woodruff, UT Southwestern) in kinase buffer (20 mM Tris-Cl pH 7.5, 50 mM NaCl, 10 mM MgCl_2_, 1 mM DTT, 0.2 mM ATP), for 1 hour at 23°C. Phosphorylated GST::BUB-1 was then directly added to CDC-20::3xFLAG bound to beads to a final GST::BUB-1 concentration of 100 nM. Binding was performed for 2 hours at 4°C, after which beads were washed with lysis buffer 4 times. Proteins were eluted from the beads by resuspending in SDS sample buffer before analysis by immunoblotting.

### Preparation of worm lysates

L4-stage worms were injected with dsRNA at a concentration of 1 μg/μL and incubated at 20°C for 36-48 hours. 60 worms per condition were washed with M9 plus 0.1% Tween 20, resuspended in SDS-sample buffer and lysed by sonication followed by boiling.

### Immunoblotting (Antibody List Table S5)

For binding assays, samples were run on 4-15% gradient Mini-Protean gels (BioRad) and transferred to PVDF membranes using a TransBlot Turbo system (BioRad). Membranes were blocked for 1 hour in TBS-Tween 0.05% (TBS-T) with 5% skim milk and antibodies were incubated in either the same buffer or in 5% BSA. Membranes were developed using Super Signal West Femto Max Sensitivity Substrate (ThermoFisher) before imaging on ChemiDoc system (BioRad). For RNAi blots, the equivalent of 5-10 adult worms was loaded into 4-12% NuPAGE Bis-Tris gels (Invitrogen). Proteins were transferred overnight to nitrocellulose membranes, after which membranes were blocked and antibodies incubated in TBS-T with 5% milk. Membranes were developed using Western Bright Sirius substrate (Advansta) before imaging on ChemiDoc System (BioRad).

### Screen for Putative PLK-1 sites (Table S6)

Putative PLK-1 phosphorylation sites in *C. elegans* BUB-1 were identified using GPS Polo 1.0 (Xue et al., 2005), with filter set to “all”. Sites were prioritized by comparing conservation with other closely related *Caenorhabditis* species, matching consensus polo motif [N/D/E/Y, X, S/T, F], and a Polo GPS score above 1.5. Sequences of *Caenorhabditis bub-1* genes were aligned using MAFFT. The full list of sites is available in Supplemental Table 6.

### AlphaFold Structure Prediction

Structure prediction for CDC-20 (residues range: 136-507) and BUB-1 (residues range: 424-458) ABBA motif interaction was performed using AlphaFold2 ColabFold notebook (Jumper et al., 2021; Mirdita et al., 2022). Structural analysis and depiction were carried out in PyMol (DeLano, 2002).

### Imaging analysis and quantifications

Microscope images were processed using ImageJ. NEBD was defined as the time point where free nuclear histone signal dissipates into the cytosol and spindle forces start pushing on chromosomes. Anaphase was the first time point where the separation of sister chromatids was evident. DNA decondensation was the time point where chromosome signals become dimmer and their area starts expanding.

Kinetochore intensities were measured by drawing a circle around the chromosome area (mCherry::H2b) and transferred to the GFP channel. After recording integrated intensity, the circle was expanded by 3 pixels to calculate local background signal.

For PLK-1 inhibition assays, kinetochore signals were quantified 60 seconds before anaphase onset. A 10 pixel wide linescan was drawn across the chromosomes and signal intensities were quantified as described before (Moyle et al., 2014). Imaging depth variations in signal intensities were corrected by dividing by the mCherry::H2b signal.

### Statistical analyses

P values were calculated in Prism software (Graphpad) using unpaired t tests. P values are displayed as follows: ns = p>0.05; * = p<0.05; ** = p<0.01; *** = p<0.001; **** = p < 0.0001. Error bars display the 95% confidence interval.

---

**Table S1:**
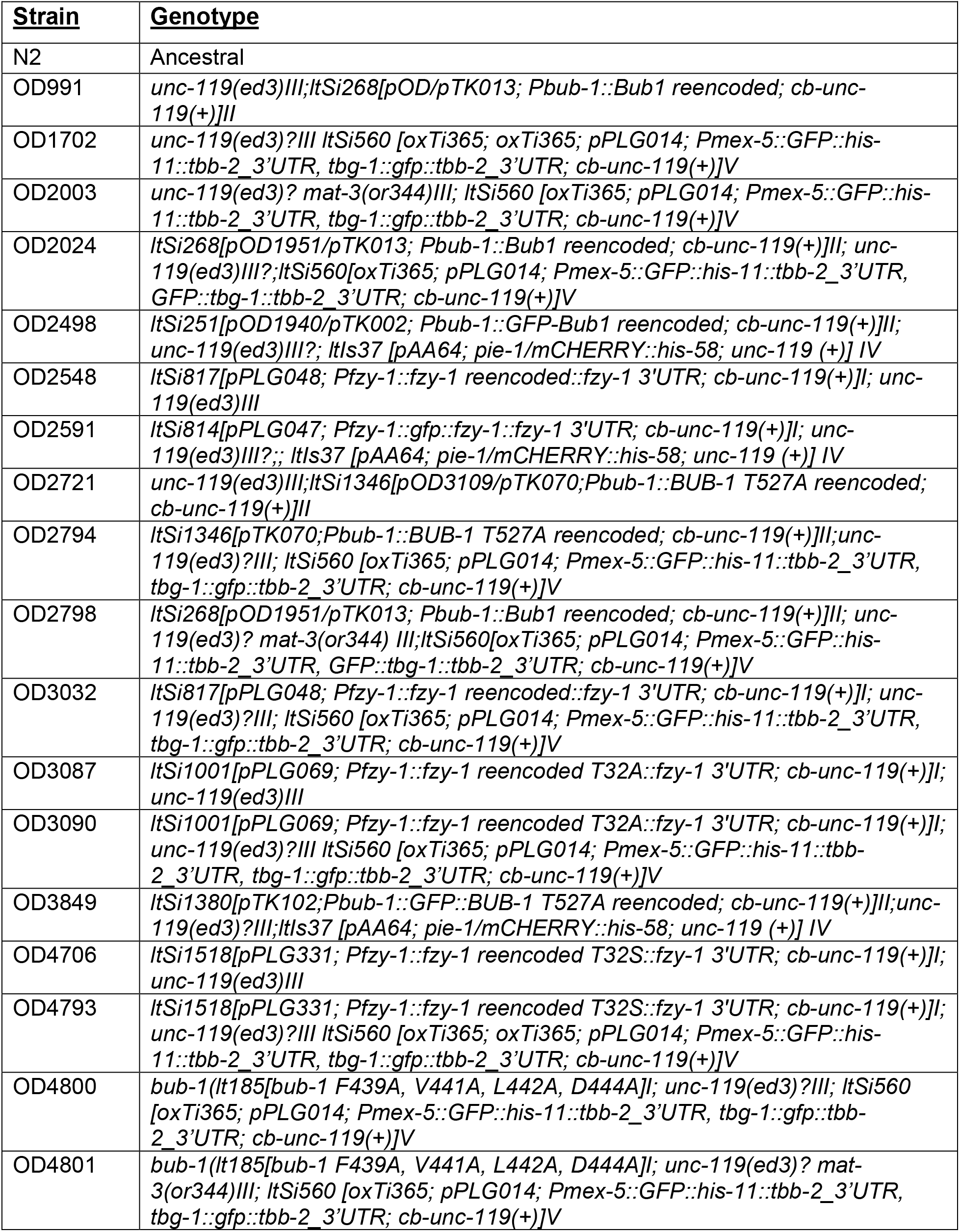
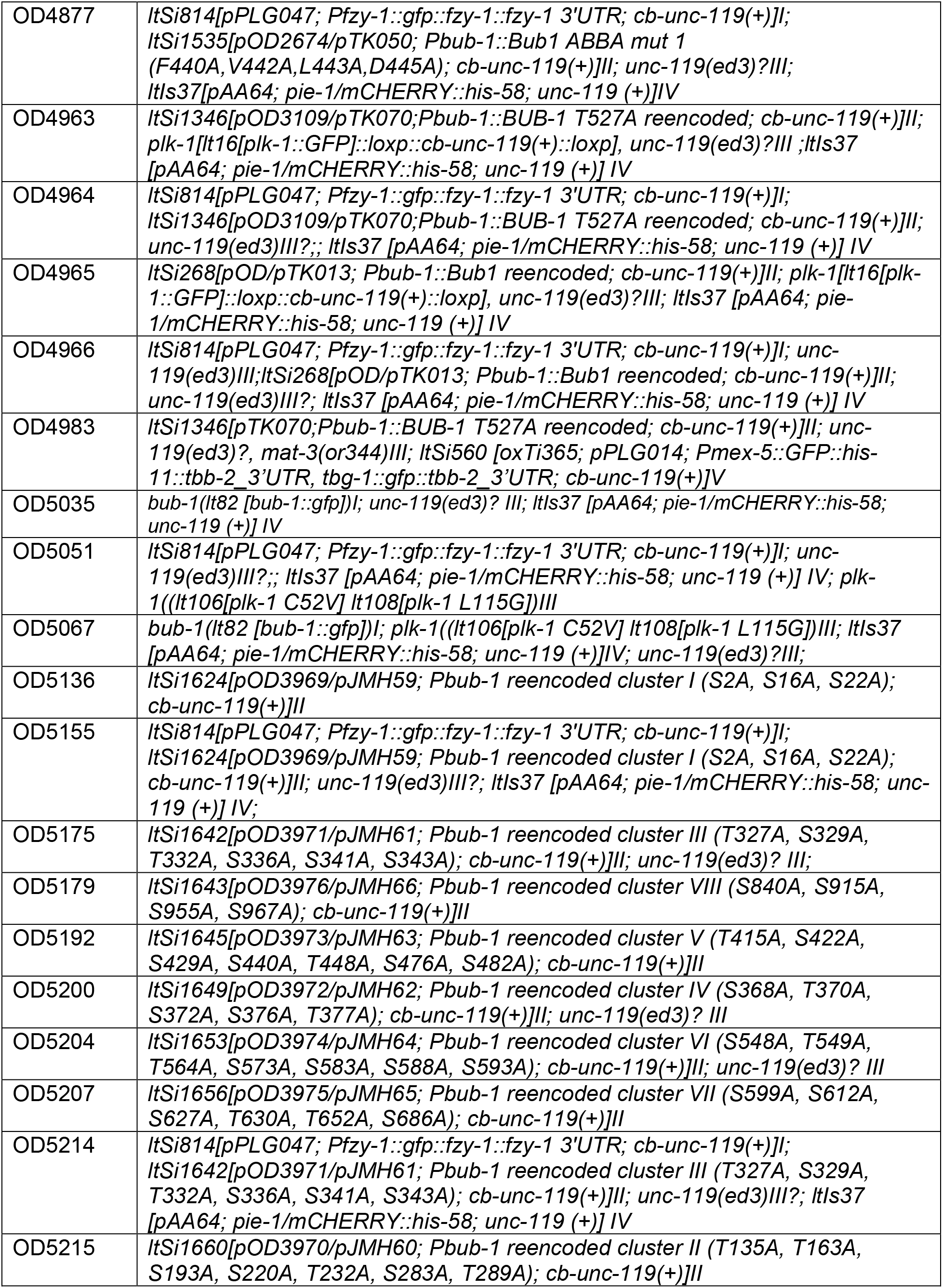
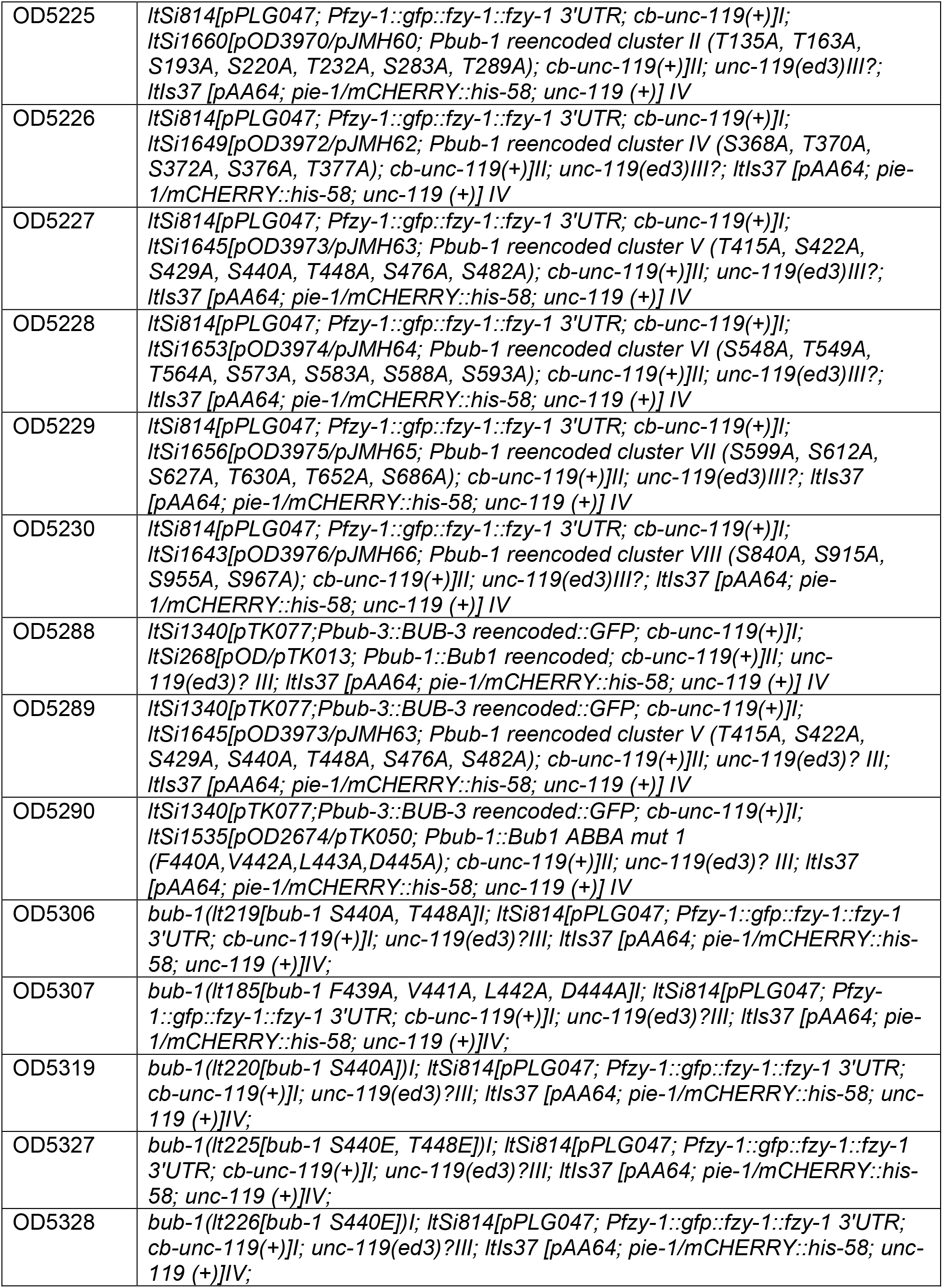
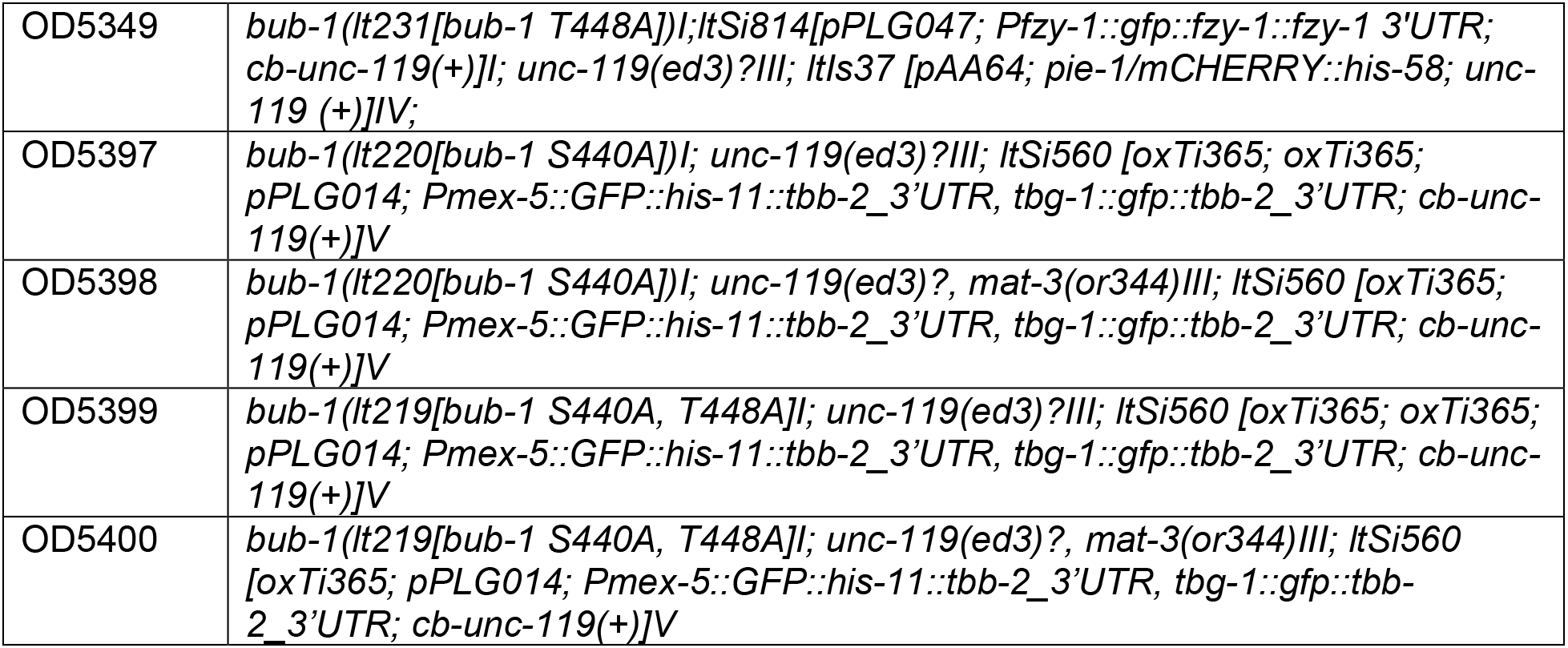
*C. elegans* strains.

**Table S2:**
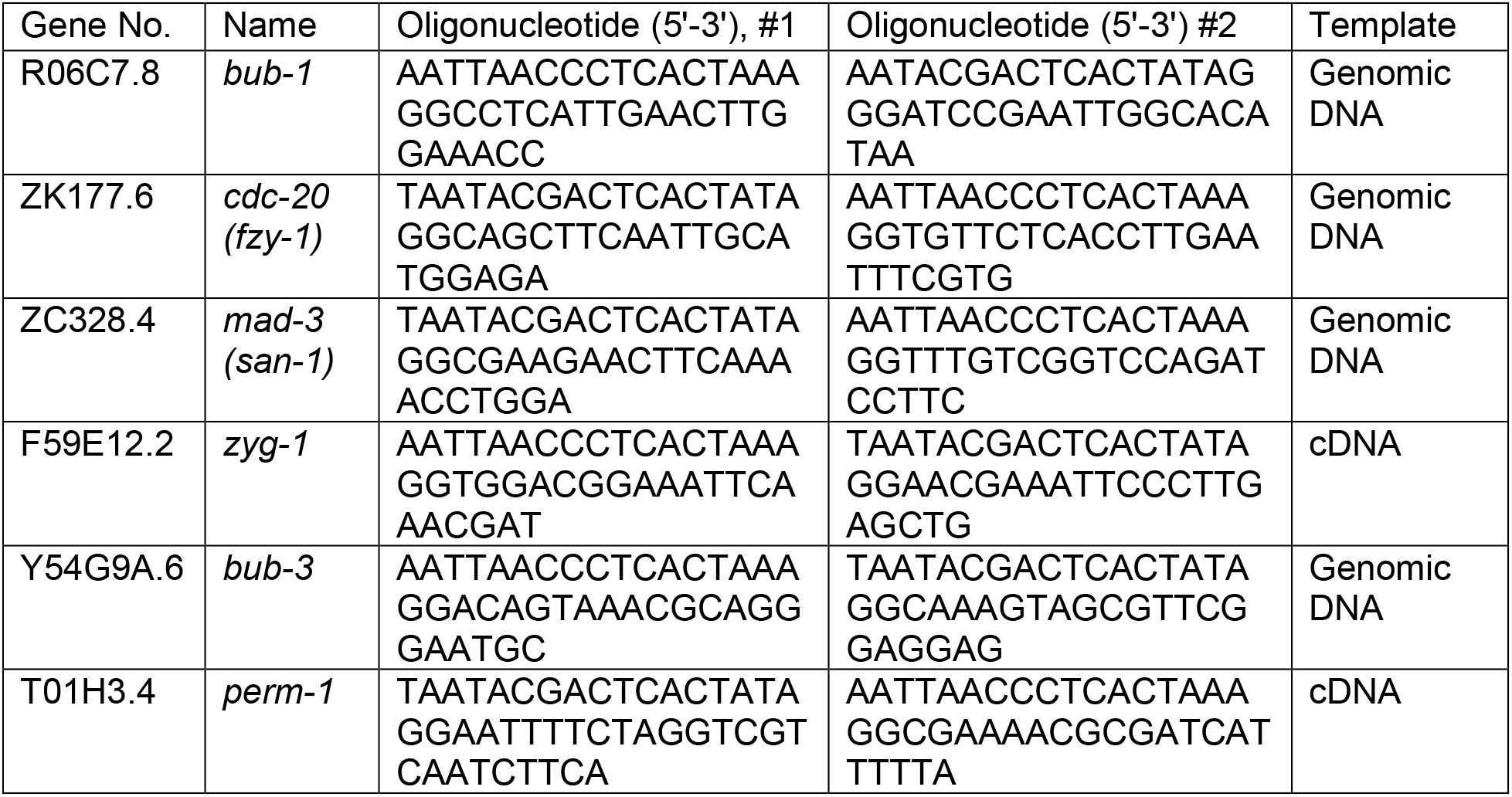
dsRNAs used in this study.

**Table S3:**
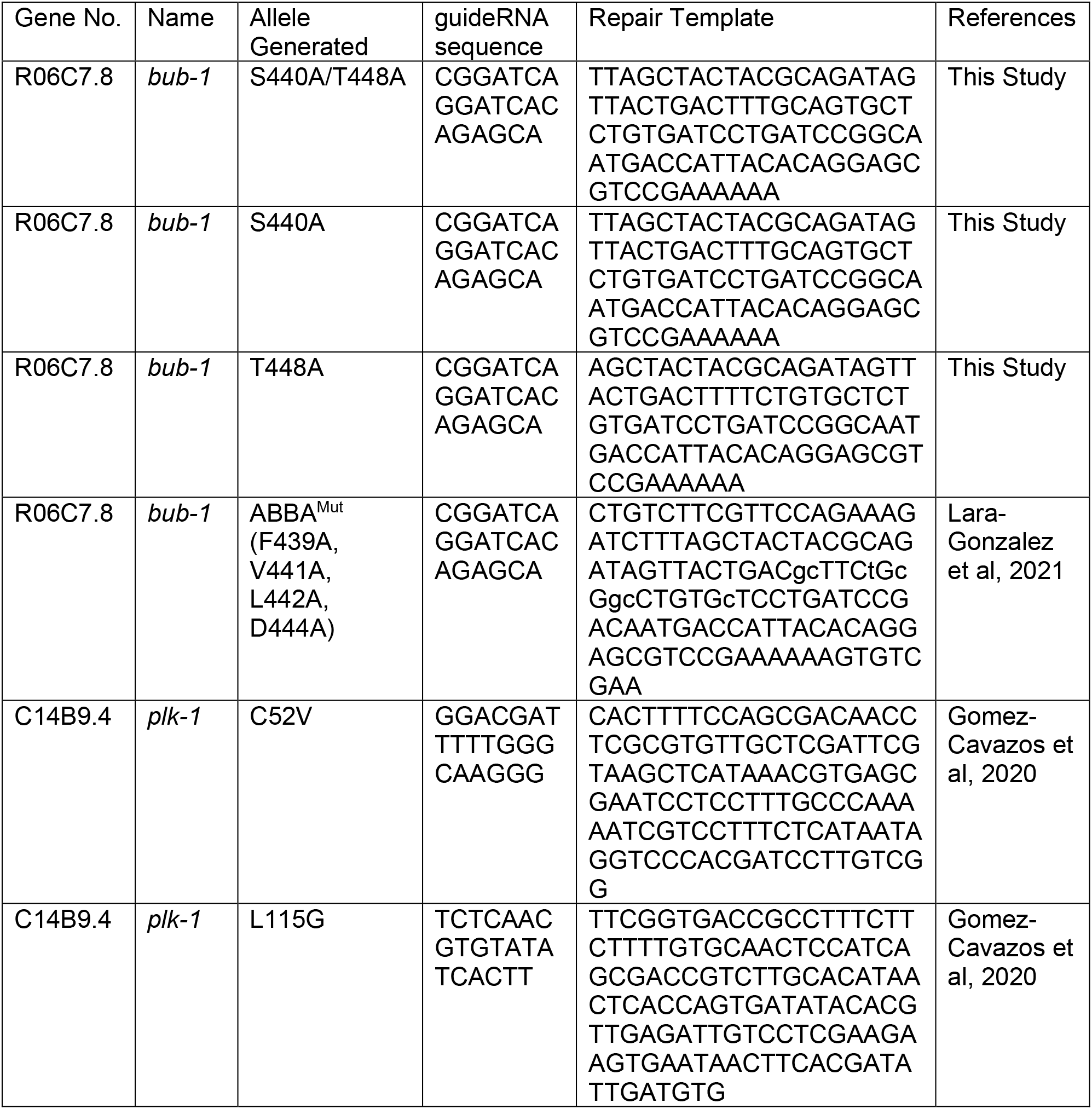
CRISPR gRNA and repair template sequences.

**Table S4:** Plasmids for protein expression:

| Plasmid # | Description | Bacterial Selection | Figures |
| --- | --- | --- | --- |
| pOD4316/pPLG352 | pCDNA 3.1(+) FZY-1::3xFLAG (C. elegans CDC-20) | Ampicillin | 3D, 4G |
| pOD4314/pPLG350 | pGEX-6P-1 GST::BUB-1 (280-472) | Ampicillin | 3D, 4G |
| pOD4374/pPLG410 | pGEX-6P-1 GST::BUB-1 (280-472) ABBA<sup>Mut</sup> | Ampicillin | 3D, 4G |
| pOD5239/pJMH75 | pGEX-6P-1 GST::BUB-1 (280-472) S440A/T448A | Ampicillin | 3D, 4G |
| pOD5236/pJMH72 | pGEX-6P-1 GST::BUB-1 (280-472) S440A | Ampicillin | 3D, 4G |

**Table S5:** Antibodies.

| Antibody Name | Target | Host Species | Stock Concentration (mg/mL) | Source |
| --- | --- | --- | --- | --- |
| OD140 | GST | Rabbit | 7.72 | Dammermann et al, 2008 |
| OD31-B | BUB-1 (aa 287-665) | Rabbit | 2.8 | Oegema et al. 2001 |
| OD220 | CDC-20 (aa 1-160) | Rabbit | 5.7 | Kim and Lara-Gonzalez et al, 2017 |
| DM1α | α-tubulin | Mouse | 1 | Sigma (T9026) |
| C4 | Actin | Mouse | 1 | Millipore (MAB1501) |
| M2 | FLAG | Mouse | 1 | Sigma (F1804) |
|  | Anti-Rabbit IgG, HRP conjugated | Goat | 0.8 | Jackson ImmunoResearch (111-035-003) |
|  | Anti-Mouse IgG, HRP conjugated | Donkey | 0.8 | Jackson ImmunoResearch (715-035-150) |
|  | Anti-Mouse Light Chain Specific, HRP conjugated | Goat | 0.8 | Jackson ImmunoResearch (115-035-174) |
|  | Anti-Rabbit IgG, HRP conjugated | Donkey | 0.8 | Jackson ImmunoResearch (711-035-152) |
|  | Mouse IgG, AP conjugated | Donkey | 1 | Jackson ImmunoResearch (715-055-150) |

**Table S6:**
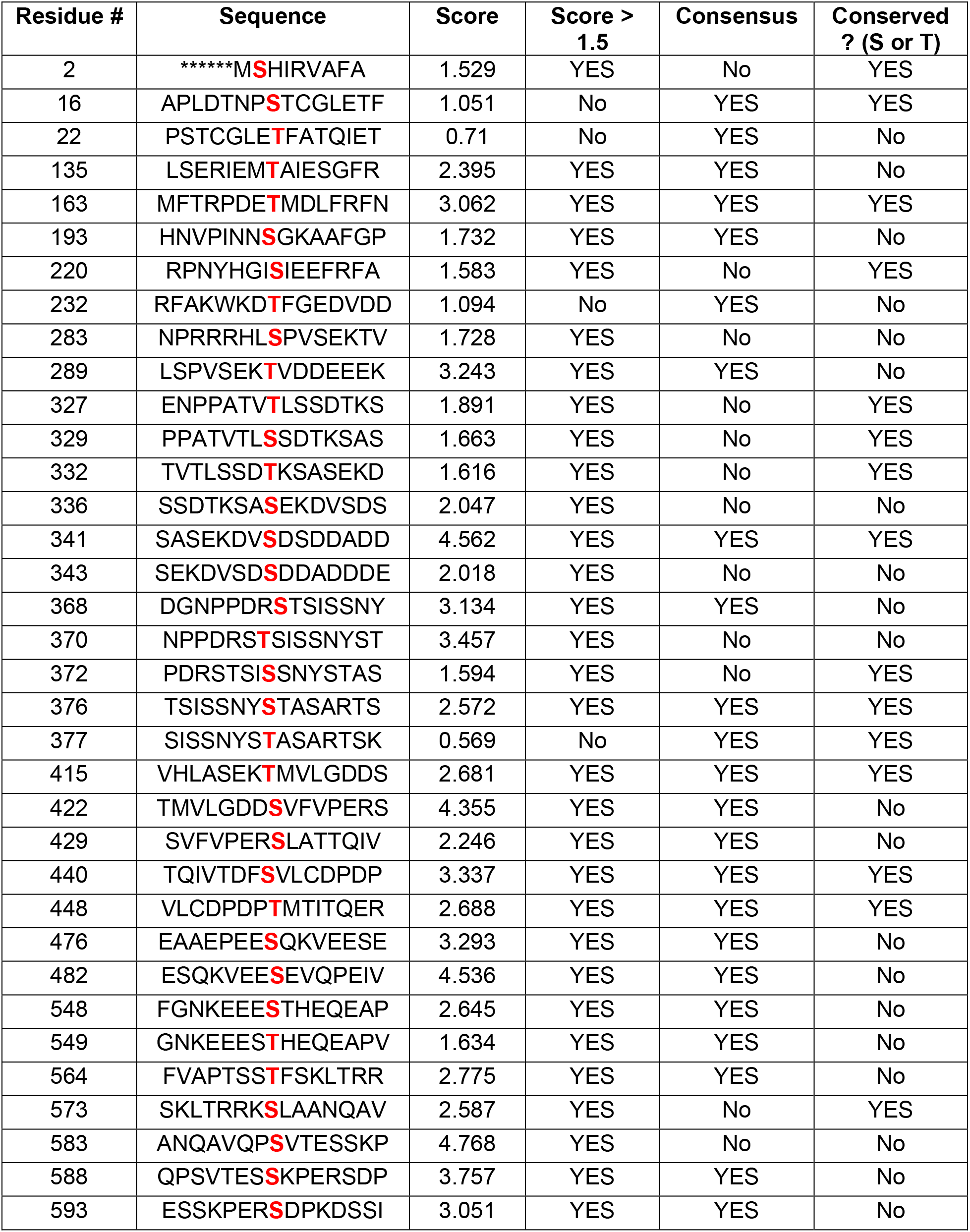
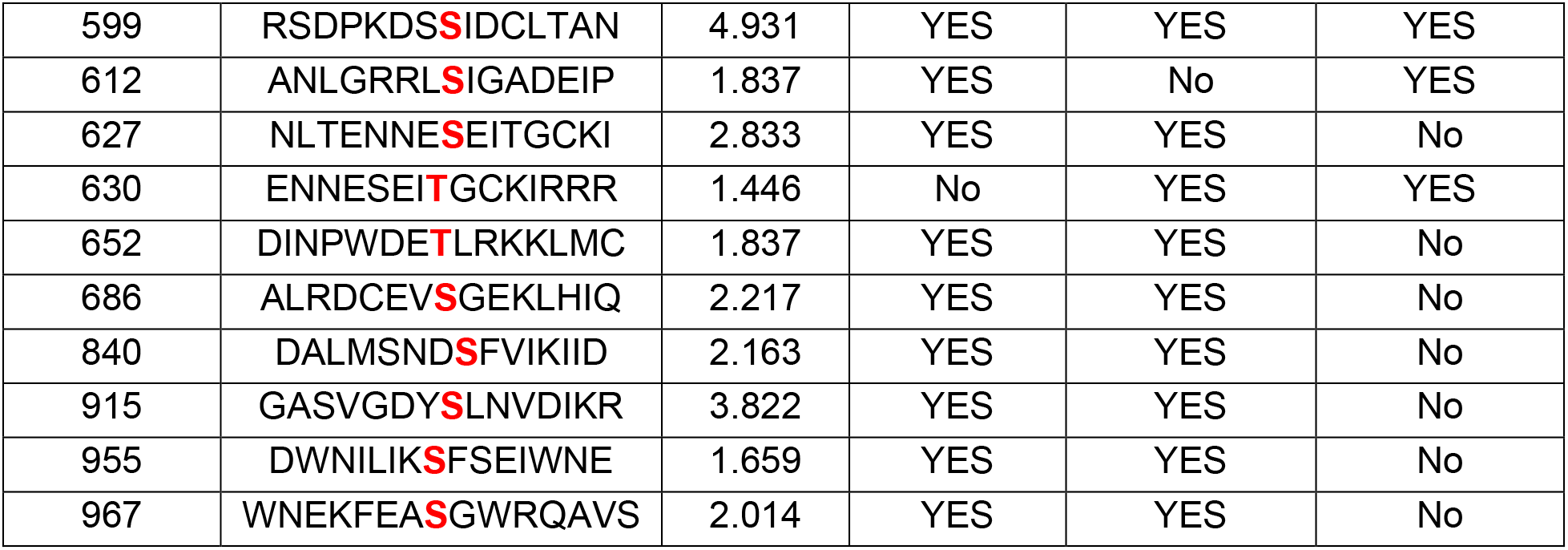
List of sites mutated in clustered mutagenesis screen (related to figure 4)

